# The emergence of E-Cadherin spot junctions between germline and somatic cells facilitates late stage oogenesis in *Drosophila*

**DOI:** 10.1101/2025.01.27.635036

**Authors:** Vanessa Weichselberger, Ramya Balaji, Marta Rodriguez-Franco, Anne-Kathrin Classen

**Affiliations:** EMBL Barcelona, 08003 Barcelona, Spain; Aix Marseille University, CNRS, UMR 7288, IBDM, 13288 Marseille, France; Hilde-Mangold-Haus, University of Freiburg, Freiburg, Germany; Faculty of Biology, University of Freiburg, Freiburg, Germany; Spemann Graduate School of Biology and Medicine (SGBM), University of Freiburg, Freiburg, Germany; Signalling Research Centers BIOSS and CIBSS, University of Freiburg, Freiburg, Germany

## Abstract

Coordinated tissue morphogenesis relies on precise interactions between distinct cell lineages. During development of the *Drosophila* oocyte, interactions between different somatic follicle cell types and the germline-derived nurse cells and the future oocyte are crucial for coordinating oogenesis. However, the molecular basis of their interaction throughout enormous morphological changes remains largely unexplored. During late oogenesis, nurse cells transfer their cytoplasmic content into the oocyte, while anterior follicle cells (AFCs) must closely envelop the nurse cells to induce phagoptosis and remove their remnants. We identify E-cadherin-based spot junctions as novel adhesive complexes at this essential soma-germline interface. AFCs reorganize their E-cadherin from intraepithelial adherens junctions to their apical surface to form interlineage spot junctions with nurse cells, representing a switch in adhesive and thus mechanical coupling. The distribution and density of these junctions is stably organized, even as the interface undergoes dramatic size changes. Additionally, we show that these junctions integrate into a microvilli-rich nurse cell surface, influencing microvilli architecture. Importantly, disruption of E-cadherin spot junctions impairs nurse cell enveloping. Our findings reveal a switch from intra-lineage to inter-lineage mechanical coupling that is driven by the formation of E-Cadherin spot junctions and highlights their essential role in ensuring robust soma-germline coupling and coordination during nurse cell dumping in late oogenesis.

## INTRODUCTION

Oogenesis is a developmental process highly dependent on the coordination of two distinct lineages, namely the germline and the surrounding somatic follicle cells. Across species from invertebrates to vertebrates, oocytes develop within a germline cyst that comprises the oocyte and nurse cells. Nurse cells play a critical role by supplying the oocyte with essential materials and facilitating its growth. However, to produce a single mature oocyte, nurse cells must ultimately be removed. This removal relies on the interaction with surrounding follicle cells. A group of follicle cells undergo specialized differentiation to eventually envelop and remove nurse cells through phagoptosis. Crucially, to allow this critical step to take place follicle cells and nurse cells have to closely interact to ensure coordination during massive changes in morphology. This dynamic cooperation underscores the intricate interdependence between nurse cells and follicle cells during oogenesis.

During gametogenesis, germline cells and surrounding somatic cells must coordinate their shared development to create fertile gametes. During mouse oogenesis, germline-derived cells reveal a remarkable specialization as oocytes and nurse cells, the later assisting the growth of the future oocyte. At the end of primordial oogenesis shortly after birth, nurse cells are eliminated and only oocytes survive (Lei and Spradling 2016, Doherty, Amargant et al. 2022, Niu and Spradling 2022, Spradling, Niu et al. 2022). The cellular mechanisms of how nurse cell specialization is achieved, how their growth and function is supported by the surrounding somatic cells and how the nurse cells are efficiently and timely eliminated are currently not understood.

During *Drosophila* oogenesis, one oocyte develops in a cyst connected to 15 nurse cells. The somatic follicle cells surround the cyst and closely coordinate their differentiation, growth and morphogenesis with that of nurse cells and the oocyte. In early oogenesis (stage 6), the follicle epithelium differentiates into posterior, main body, centripetal, anterior and border cells by a well-described combinatorial action of Notch, Jak-Stat, and EGFR signaling (Dobens and Raftery 2000, Horne-Badovinac and Bilder 2005, Roth and Lynch 2009). In mid-oogenesis (stage 9), follicle cells rearrange over the germline cyst to build precisely matching interlineage units. Specifically, posterior and main body cells make contact with the growing oocyte, whereas anterior cells spread to completely cover all nurse cells (Dobens and Raftery 2000, Horne-Badovinac and Bilder 2005, Grammont 2007, Kolahi, White et al. 2009, Roth and Lynch 2009, Chlasta, Milani et al. 2017, Balaji, Weichselberger et al. 2019, Lamire, Milani et al. 2020, Weichselberger, Dondl et al. 2022). This cell-type specific distribution of follicle cell fates over oocyte and nurse cells into matching interlineage compartments needs to be maintained and stabilized in late oogenesis as it is essential for the successful development of a fertile oocyte: Posterior and main body follicle cells enveloping the growing oocyte will help build the future chorion and differentiate egg shell structures (Mauzy-Melitz and Waring 2003, Duhart, Parsons et al. 2017). In contrast, anterior follicle cells (AFCs) and centripedal cells ensure the continuous coverage of nurse cells as nurse cells dramatically change size during egg chamber morphogenesis. Specifically, nurse cells massively grow and increase in volume and surface area during stage 10 and rapidly decrease their volume during stage 11 in a process called nurse cell dumping. Nurse cell dumping is the process in which nurse cells push their entire cytoplasmic content into the oocyte (Huelsmann, Ylanne et al. 2013, Timmons, Mondragon et al. 2016, Imran Alsous, Romeo et al. 2021, Logan, Chou et al. 2022, Niu and Spradling 2022, Spradling, Niu et al. 2022, Giedt and Tootle 2023). After the transfer of all cytoplasmic content into the oocyte, nurse cells have fulfilled their role in oogenesis and therefore must be removed. The removal is executed by the AFCs, which during dumping envelope each nurse cell individually, induce phagoptosis and eventually engulf all nurse cell remnants (Etchegaray, Timmons et al. 2012, Peterson and McCall 2013, Timmons, Mondragon et al. 2016, Lebo and McCall 2021). This sequence of processes supplies the oocyte with critical factors for early embryogenesis and ensures the reduction of the germline cyst to a single oocyte.

It is conceivable, that anterior follicle cells must maintain intimate contact with nurse cells to ensure complete nurse cell coverage during the dramatic growth and shrinkage phases in late oogenesis. Moreover, during dumping nurse cells display major actomyosin contraction, which poses an additional challenge for a stable interface between AFCs and nurse cells (Wheatley, Kulkarni et al. 1995, Ferreira, Prudencio et al. 2014, Imran Alsous, Romeo et al. 2021). We previously described how AFCs cover the outward facing nurse cell surface prior to stage 10A in an affinity-dependent spreading process regulated by the transcriptional co-regulator Eyes absent (Eya) (Weichselberger, Dondl et al. 2022). Yet, it is not understood how subsequently AFC and nurse cells interact to stabilize and coordinate their interface during volume changes in late oogenesis and rapid cortical contractions of nurse cells during dumping. Thus far, the role of the interface between the somatic cell surface and the germline has been little studied, despite evidence that this interface harbors signaling, mechanical, and nutritional roles during oogenesis (Waksmonski and Woodruff 2002, Bohrmann and Zimmermann 2008, Roth and Lynch 2009, Meehan, Kleinsorge et al. 2015, Balaji, Weichselberger et al. 2019, Isasti-Sanchez, Munz-Zeise et al. 2021, Row, Huang et al. 2021, Weichselberger, Dondl et al. 2022, Giedt, Thomalla et al. 2023, Mallart, Netter et al. 2024).

Adhesion between AFCs and nurse cell surfaces could provide one possible solution to stabilize the interaction between these two cell lineages. Previous work has revealed that during early oogenesis adhesion mediated by homophilic E-cadherin interactions between germline cells and follicle cells ensures the correct positioning of the oocyte at the posterior pole of the early egg chamber (Godt and Tepass 1998, Gonzalez-Reyes and St Johnston 1998, Riparbelli, Persico et al. 2022). However, adhesive structures relevant for late oogenesis and the stabilization of soma-germline interactions have not been elucidated. Importantly, somatic follicle cells are generally considered to represent an epithelial tissue type with classical cell-cell adhesion mediated via E-cadherin dependent adherens junctions and a defined apical-basal polarity (Horne-Badovinac and Bilder 2005, Duhart, Parsons et al. 2017). Notably, the apical surface of follicle cells contacts the germline, while the basal surface faces outward and is covered by a basement membrane (Isabella and Horne-Badovinac 2016, Osswald, Barros-Carvalho et al. 2022). This arrangement is highly unusual, as apical epithelial surfaces typically face an open luminal environment. How apical diversification arises to enable its interaction with a different cell type remains largely unexplored. In this study, we characterize the emergence and function of E-cadherin–based spot junctions that form during late *Drosophila* oogenesis, where they stabilize soma–germline contacts to produce fertile eggs.

## RESULTS

### E-cadherin redistributes into clusters at the apical follicle cell surface

We wanted to investigate the cell biological basis and function of the contact between nurse cells and AFCs during late oogenesis (stage 10A and onwards). The precise spatial matching of contact between nurse cells and AFCs is established during mid-oogenesis (stage 9) when AFCs spread over the nurse cell compartment **(Fig.1A)** (Weichselberger, Dondl et al. 2022). We found that during AFC spreading, intra-epithelial adherens junctions (AJ) in AFCs marked by staining for E-cadherin (E-cad), encoded by the Drosophila *shotgun (shg)*, break and largely dissolve **(Fig.1B,C)**, consistent with a previous report on AFC flattening (Grammont 2007). However, we also observed that the loss of E-cadherin at intra-epithelial AJs coincided with the appearance of E-cadherin clusters in the apical surface of AFCs where they spread over the nurse cell surface **(Fig. 1B,C)**. In contrast, E-cadherin was retained in belt-like AJs in posterior and main body cells, and the apical surfaces of posterior and main body cells did not contain E-cadherin clusters when still in contact with nurse cells, or later with the oocyte **(Fig.1B)**. Thus, during the process of AFC spreading, E-cadherin is redistributed from intra-epithelial interfaces in AFCs to their apical surface in contact with nurse cell surfaces **(Fig. 1D)**.

**Figure 1.**
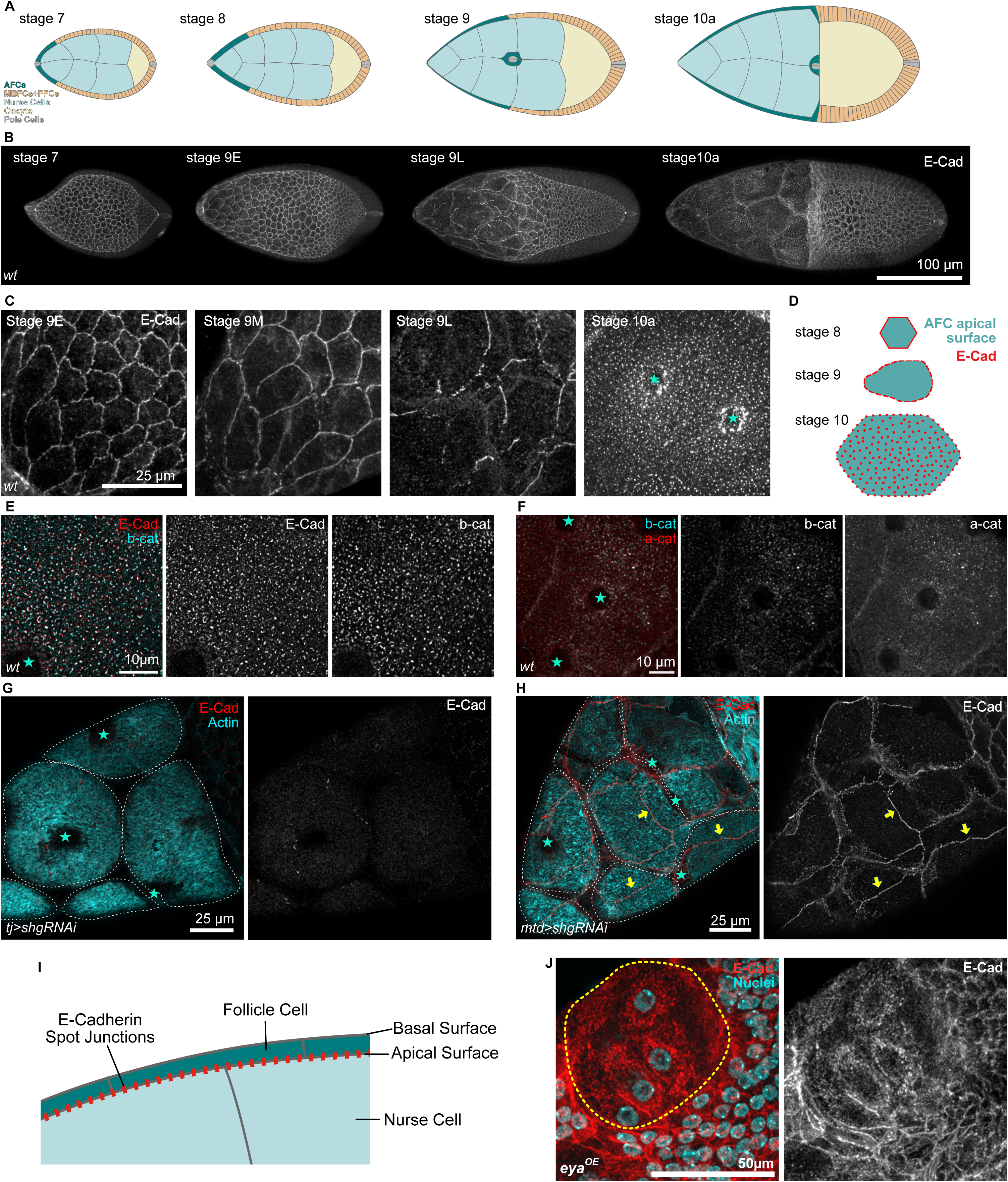
Follicle cell adherens junctions are remodelled into germline-soma spot junctions. **A** Illustrations of *Drosophila* egg chambers during mid oogenesis from stages 7 to 10A. **B** Max. projections of confocal images of *Drosophila* egg chambers of di=erent developmental stages stained for E-Cad. Note changes in E-cadherin organization over time, specifically in the anterior egg chamber. **C** Max. projection of confocal images depicting AFCs at di=erent developmental stages stained for E-Cadherin. Note fragmentation of adherens junctions in stage 9 and establishment of E-cadherin clusters on apical cell surface by stage 10A. **D** Illustration of the apical surface and E-cadherin organization of AFCs in di=erent developmental stages. **E** Airyscan image depicting AFCs of a stage 10A egg chamber stained for E-cadherin and β-catenin. Cyan stars mark position of AFC nuclei. **F** Max. projection of confocal images depicting AFCs of a stage 10A egg chamber stained for β-catenin and α-catenin. Cyan stars mark position of AFC nuclei. **G** Max. projection of confocal images depicting a stage 10A egg chamber, expressing *shg-RNAi* under the control of TJ-Gal4 in follicle cells, and stained for F-Actin and E-Cadherin. Cyan stars mark location of AFC nuclei. **H** Max. projection of confocal images depicting a stage 10A egg chamber, expressing *shg-RNAi* under the control of MTD-Gal4 in germline cells, and stained for F-Actin and E-Cadherin. Cyan stars mark location of AFC nuclei. Yellow arrows mark cell-cell junctions between AFCs. Note E-cadherin presence at AFC cell-cell junctions. **I** Illustration of Nurse cell-follicle cell interface of a stage 10 egg chamber depicting E-Cad spot junction localization. **J** Max. projection of confocal images depicting a zoom into cells ectopically expressing eya and therefore increased apical surface areas in contact with nurse cells. Egg chamber shown in Fig. S1G. Stained for E-Cad and Nuclei (DAPI). Yellow dotted line encircles cells with ectopic expression of eya. Note lack of E-cadherin at follicle cell junctions and presence of E-cadherin spots on their apical surface.

From stage 10A onwards when all nurse cells were covered by AFCs, clusters of E-cadherin were present within the entire AFC-nurse cell interface. To investigate if the observed clusters may represent functional junctions, we analyzed the localization of additional AJ markers. We found that β-catenin (Armadillo, arm), α-catenin (α-cat) and Par-3 (Bazooka, Baz) localized in E-cadherin clusters **(Fig. 1E,F, Fig.S1A)**. These observations suggest that E-cadherin clusters at the AFC-nurse cell interface colocalize with bona fide AJ proteins. In contrast, regulators of apical-basal polarity, such as Crumbs (Crbs) and atypical Protein Kinase C (aPKC) did not localize with E-cadherin clusters, neither did other adhesion proteins like N-cadherin and Echinoid (Ed) **(Fig. S1B-E)**. Combined, these observations show that E-cadherin is removed from intra-epithelial adherens junctions during AFC spreading and reorganizes into clusters in the apical follicle cell surface facing nurse cells from stage 10A onwards.

### E-cadherin forms functional spot junctions between follicle and nurse cells

Nurse cells express E-cadherin and nurse cell specific functions for E-cadherin have been described (Huelsmann, Ylanne et al. 2013, Loyer, Kolotuev et al. 2015). To investigate if the E-cadherin clusters in the follicle cell membrane may in fact represent homophilic adhesion between E-cadherin on follicle and nurse cell surfaces, we knocked down E-cadherin expression in follicle cells or in nurse cells. We employed RNAi-mediated knockdowns under the control of either TJ-Gal4 or mat-α-tub-Gal4, respectively, which strongly reduced the levels of E-cadherin in the targeted cell types, but allowed egg chambers to develop to stage 10A. Consistent with the hypothesis that E-cadherin clusters represent functional adhesive junctions between follicle and nurse cells, we find that knock-down of E-cadherin in either cell type alone is sufficient to abolish the occurrence of E-cadherin clusters at the follicle-nurse cell interface from stage 10A onwards **(Fig. 1G,H)**. We therefore conclude that E-cadherin clusters are functional adhesive spot junctions between the apical follicle cell surface and the nurse cell surface **(Fig. 1I)**.

Interestingly, knock-down of E-cadherin in the germline not only prevented E-cadherin cluster formation at the follicle-nurse cell interface but also, in a non-cell-autonomous manner, lead to localization of E-cadherin to intraepithelial junctions between AFC at stage 10 **(Fig.1H)**. This suggests that E-cadherin in nurse cells may act as a sink for E-Cadherin in AFCs. The contact surface between AFCs and nurse cells is significantly larger than the contact surface between AFCs themselves at stage 10. Therefore, the probability to form E-Cadherin trans bonds is biased towards the AFC-nurse cell interface. By removing the trans-binding partner of E-cadherin in the apical surface through E-cadherin knockdown in the germline E-cadherin trans bond formation is biased towards follicle cell interfaces. To further investigate this hypothesis, we ectopically increased the apical surface in MBFCs, that normally do not increase their apical surface and transition onto the oocyte, to test whether a large apical surface shared with nurse cells is sufficient to induce E-Cadherin cluster formation at the FC-nurse cell interface. Ectopic expression of Eya in MBFCs causes their ectopic spreading by increasing their apical surface area in contact with nurse cells (Weichselberger, Dondl et al. 2022). We found that ectopic expression of Eya in cell clones and the resulting spreading of MBFCs on nurse cell surfaces was sufficient to induce the ectopic formation of apical spot junctions during stage 9 and 10A **(Fig. 1J, Fig. S1F)**. These results indicate that increasing the probability of trans-bond engagement on the expanding apical surface of AFCs can bias E-cadherin localization and spot junction formation toward the nurse cell interface.

A closely related molecule that may interact with E-cadherin at spot junctions is N-cadherin. Follicle cells express N-cadherin and have been previously shown to upregulate N-cadherin to compensate for a loss of E-cadherin in early stages of egg chamber development (Grammont 2007). We therefore analyzed N-cadherin localization in E-cadherin mutant clones during stage 10A to understand if N-cadherin may initially contribute to the formation or even compensate for loss of E-cadherin in spot junctions. In wild type egg chambers at stage 10A, N-cadherin localized to intra-epithelial adherens junctions but not to the follicle-nurse cell interface. Upon E-cadherin knock-down in follicle cells, increased levels of N-cadherin are present at intra-epithelial AJs at stage 10A, but N-cadherin failed to form clusters or spot junctions at the follicle-nurse cell interface **(Fig. S1G)**. This is consistent with the observation that nurse cells do not express N-cadherin and therefore cannot provide N-cadherin at their surfaces for homophilic trans-bond engagement (Loyer, Kolotuev et al. 2015). Combined our results demonstrate that E-cadherin forms functional spot junctions between the apical surface of follicle cells and nurse cells by stage 10A and that their formation is strictly dependent on homophilic trans-bond engagement between the two cell lineages.

### Spot junction organization and density remains constant during nurse cell growth

To better characterize E-cadherin spot junctions between these two cell types **(Fig. 2A)**, we first analyzed the distribution of E-cadherin spot junctions over the follicle-nurse cell interface. We analyzed the distribution of spot junctions by quantifying the nearest neighbor distance (NND). We found that that the average NND was around 1µm, which was 7% larger than in randomized distributions **(Fig.2B, Fig.S2A)** and that the minimal NND was significantly larger than in randomized samples **(Fig. S2B)**. Furthermore we found a significantly lower coefficient of variance in NNDs when compared to randomized distributions **(Fig. 2C)**. Remarkably, the NND, the CV of NNDs, the density of spot junctions and the area proportion that spot junctions made up was stable over the course of a massive follicle-nurse cell interface increase from stage 10A to stage 11 **(Fig. 2D-H, Fig. S2C).** This suggests a regulated organization of E-Cadherin spot junctions at the AFC-nurse cell interface, where spot junctions are evenly spread out over the interface with an average NND of 1µm and a stable density.

**Figure 2.**
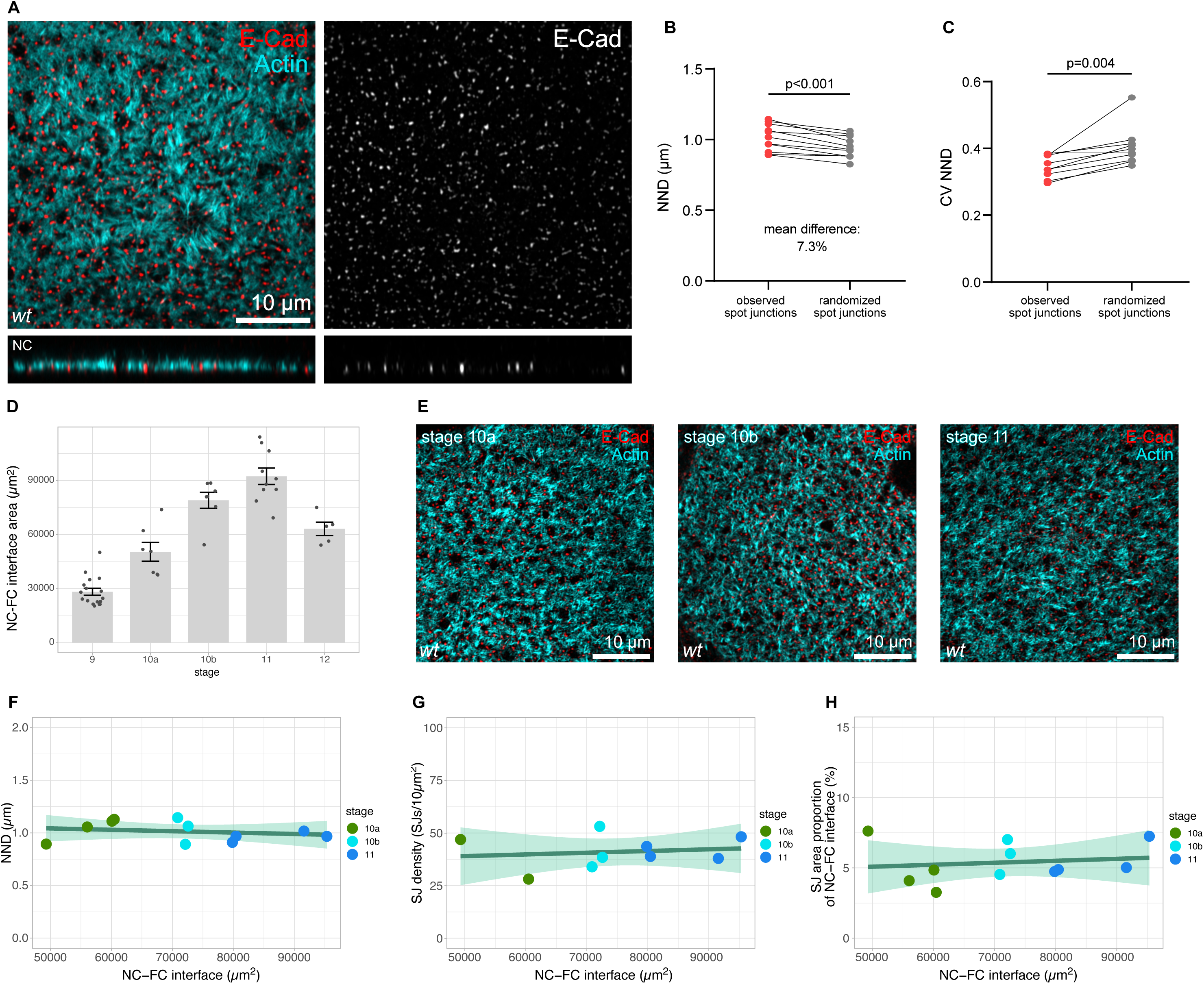
Spot junctions distribute with constant density at the follicle-nurse cell interface during growth. **A** Airyscan imaging of the AFC-nurse cell interface of a wt stage 10A egg chamber stained for E-cadherin and F-Actin. **B** Two-tailed paired t-test of the Nearest Neighbor Distance (NND) between E-cadherin spot junctions. Observed spot junction organization vs randomized organization (see Fig. S2 & Methods). n=11 egg chambers. **C** Two-tailed paired t-test of the Coe=icient of Variance (CV) of NNDs between E-cadherin spot junctions. Observed spot junction organization vs randomized organization (see Fig. S2 & Methods). n=11 egg chambers. **D** Quantification of nurse cell-to-follicle cell interface sizes at di=erent wt egg chamber stages. N egg chambers: (stage9=17, stage 10A=7, stage 10B=6, stage 11=10, stage 12=5). **E** Airyscan images of the AFC-nurse cell interface of wt egg chambers from stage 10A, 10B and 11, stained for E-cadherin and F-Actin. **F** NND between spot junctions as a function of nurse cell-to-follicle cell interface size of wt egg chambers. Linear fit with a 95% CI area. n=11 egg chambers. **G** Spot junction density as a function of nurse cell-to-follicle cell interface size of wt egg chambers. Linear fit with a 95% CI area. n=9 egg chambers. **H** Spot junction area proportion of nurse cell-to-follicle cell interface as a function of nurse cell-to-follicle cell interface size in wt egg chambers. Linear fit with a 95% CI area. n=11 egg chambers.

To better understand how spot junction distribution may be organized at the follicle-nurse cell interface, we investigated how spot junctions are integrated into the follicle cell and nurse cell actin structures. The total F-Actin staining revealed a dense actin meshwork at the AFC-NC interface **(Fig. 3A,B)**. To better characterize this structure we wanted to first distinguish follicle and nurse cell contributions to the total F-Actin staining and thus performed cell-type specific labelling experiments of F-Actin. While follicle-cell-specific expression of the actin binder utrABD-GFP failed to label this pronounced actin meshwork and rather revealed a meshwork of fine actin filaments **(Fig. 3A)**, utrABD-GFP expression in the germline revealed a complex structure resembling that of total F-actin staining **(Fig. 3B)**. Specifically, the nurse cell surface contained densely packed actin microfilaments and small actin free regions. Importantly, E-cadherin spot junctions localized to the edges of actin-free spaces rather than to actin rich regions **(Fig. 3C)**. This correlation was confirmed by the quantification of actin intensities at E-cadherin spot junctions, which was significantly lower compared to actin intensities at spot junctions with randomized distributions **(Fig. 3D)**. This suggests that the dense actin meshwork of nurse cells and E-cadherin spot junctions are organized in respect to each other. To better understand the ultrastructural architecture of these actin structures, we performed a TEM analysis of the nurse cell-follicle cell interface at stage 10A. This revealed that the surface of nurse cells was covered by microvilli, likely representing the observed dense filamentous actin meshwork **(Fig. 3E)**. Microvilli extended into the space between nurse cells and follicle cells and were absent in regions where nurse cells and follicle cells formed membrane contacts, potentially representing E-cadherin spot junctions **(Fig. 3E)**. In egg chambers expressing E-cadherin RNAi in either the follicle cells (FCs) or the germline, the membrane contacts observed in wild type egg chambers were absent **(Fig. 3F,G)**. Importantly, microvilli still formed, yet, the distance between nurse cells and follicle cells was smaller in RNAi-expressing egg chambers, with microvilli being aligned more parallel to the nurse cell surface rather than extending away from the surface **(Fig. 3E-G)**. The role of these nurse cell microvilli and their spot junction dependent architecture remains unknown as we did not observe functional effects on germline growth during late oogenesis – effects that might have been expected if disrupting microvilli architecture and therefore reduce the surface available for nutrient uptake.

**Figure 3.**
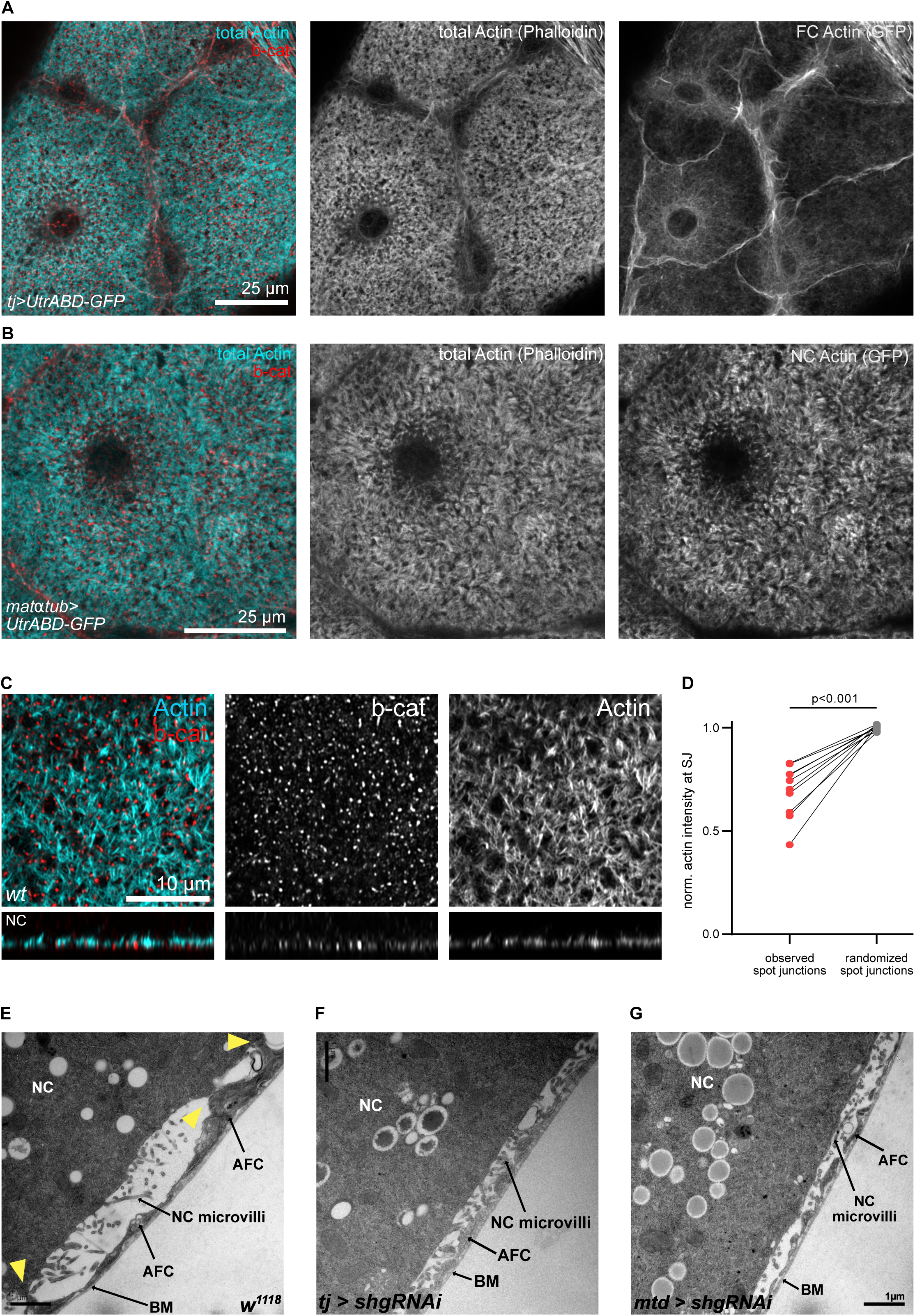
Spot junction and microvilli emergence are coordinated. **A** Max. projection of confocal images depicting AFCs in egg chamber expressing *utrABD-GFP* under the control of TJ-Gal4 in follicle cells, stained for total F-Actin (Phalloidin), GFP (follicle cell Actin) and β-catenin. **B** Max. projection of confocal images depicting AFCs in egg chamber expressing *utrABD-GFP* under the control of MTD-Gal4 in germline cells, stained for total F-Actin (Phalloidin), GFP (nurse cell Actin) and β-catenin. **C** Airyscan image of AFC-nurse cell interface of a wt stage 10B egg chamber, stained for F-Actin and β-catenin. **D** Two-tailed paired t-test of Actin intensity at spot junctions. Observed spot junction organization vs randomized organization (see Fig. S2 & Methods). n=10 egg chambers. **E** TEM image of section through AFC-nurse cell interface of a wt stage 10A egg chamber. **F** TEM image of section through AFC-nurse cell interface of a stage 10A egg chamber expressing *shg-RNAi (E-cadherin RNAi)* under the control of TJ-Gal4 in follicle cells. **G** TEM image of section through AFC-nurse cell interface of a stage 10A egg chamber expressing *shg-RNAi* under the control of MTD-Gal4 in germline cells. NC=Nurse Cell, AFC= anterior follicle cell, BM=basement membrane

### Lack of E-cadherin in follicle cells leads to ring canal blockage and nurse cell dumping failure

We next asked which function those inter-lineage E-Cadherin based spot junctions could play. Nurse cell dumping is the process in which nurse cells push all their cytoplasmic content into the oocyte **(Fig. 4A,B)**. Nurse cell dumping has been described to take place in two phases, where the transport of material from nurse cells to the oocyte is based on a surface tension based hydraulic transport in phase 1 and an active transport dependent on acto-myosin contractions of NCs in phase 2 (Imran Alsous, Romeo et al. 2021). We found that in egg chambers with follicle cell specific knock-down of E-cadherin, nurse cell dumping initiated but failed to complete **(Fig. 4C)**. While the nurse cell area proportion in mid sections of control egg chambers is 65% at the onset of nurse cell dumping, it is reduced to eventually 0% in control egg chambers, while it stagnates at 33% in egg chambers with follicle cell specific E-Cadherin knockdown **(Fig. 4D)**. Concomitantly with the failure of nurse cell dumping completion, we observed that the coordinated reduction of sizes between NC within one egg chamber was lost. Egg chambers with E-Cadherin knockdown in follicle cells had a significantly higher variance in NC sizes **(Fig. 4E, Fig. S3A)**, suggesting an impaired flow of cytoplasmic contents between germline cells. Germline cells are connected by so called ring canals that allow the flow from cell to cell. We found that in around 90% of egg chambers expressing E-cadherin RNAi, one or more ring canals were blocked **(Fig. 4F)**. Right before the onset of nurse cell dumping, nurse cells form actin cables that extend from their cell surface to the nucleus and thereby push the nucleus from the ring canals to prevent blockage **(Fig. 4G)**. It has been previously described that E-cadherin is necessary for centripetal cells to migrate between nurse cells and the oocyte and that this localization of centripetal cells is necessary for proper formation of actin cables in the contacting nurse cells (Oda, Uemura et al. 1997, Logan, Chou et al. 2022, Parsons, Mosallaei et al. 2023). We hypothesize that the lack of proper centripetal cell localization in our E-cadherin knock-down experiments causes failures in actin cable formation and as a result the blockage of ring canals specifically in the 4 nurse cells sharing a ring canal with the oocyte **(Fig. 4H)**. The blockage of a ring canal disrupts the flow and thereby changes hydraulic dynamics which can account for the drastically increased variance in nurse cell sizes.

**Figure 4.**
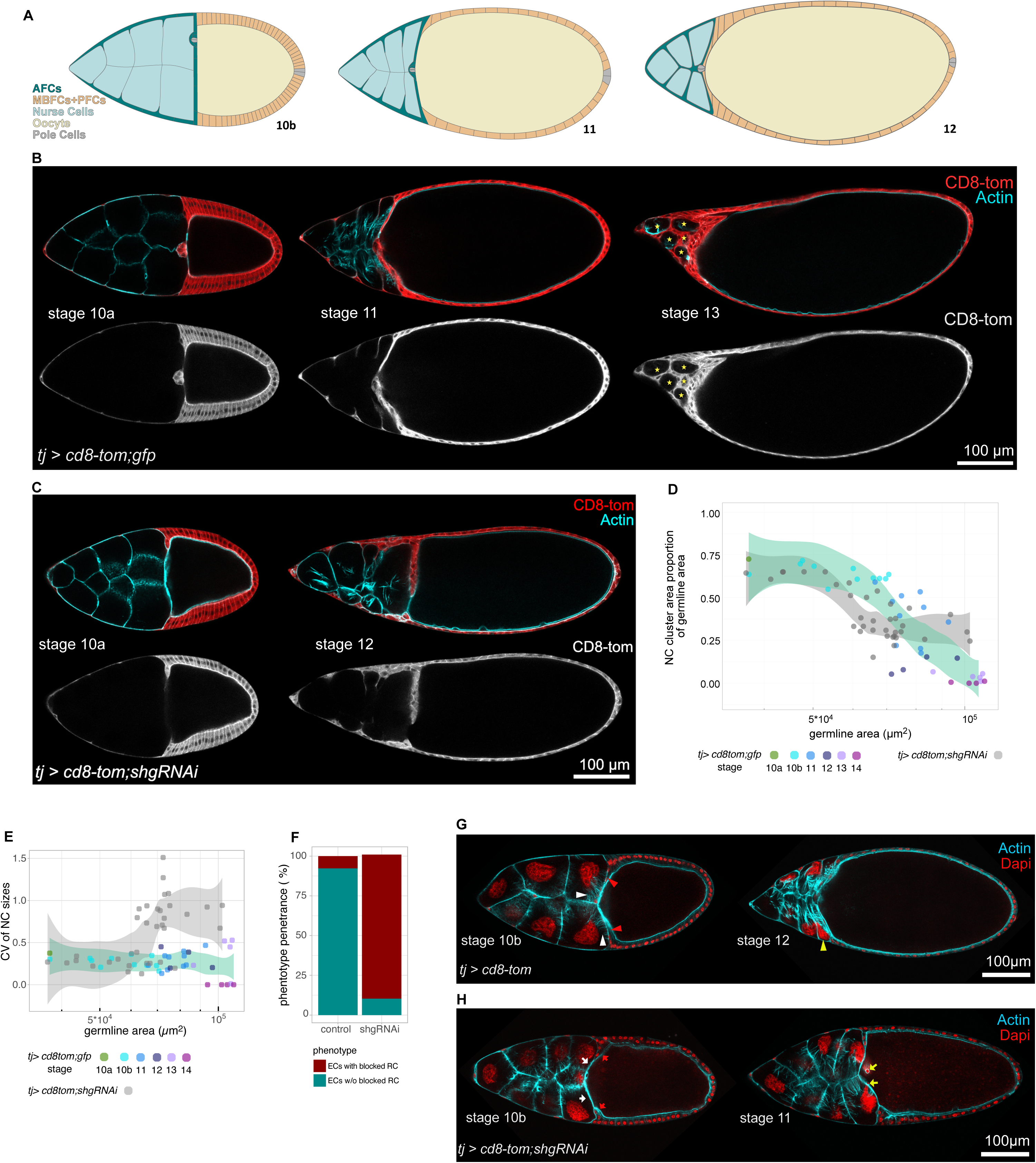
Nurse cell dumping depends on E-cadherin expression in follicle cells. **A** Illustration of *Drosophila* egg chambers undergoing nurse cell dumping. Note how AFCs completely envelop nurse cells. **B** Medial confocal sections through egg chambers from the beginning to the end of nurse cell dumping, expressing *CD8::Tomato* and *GFP* under the control of TJ-Gal4 in follicle cells, and stained for F-Actin. Yellow stars mark nurse cell remnants completely enveloped by AFCs at stage 13. **C** Medial confocal sections through chambers at stages corresponding to nurse cell dumping expressing *CD8::Tomato* and *shg-RNAi (E-cadherin RNAi)* under the control of TJ-Gal4 in follicle cells, and stained for F-Actin. **D** Quantification of nurse cell dumping progression. The proportion of the nurse cell cluster size as a function of total germline size. LOESS fitted with a 95% CI area. N egg chambers: (*tj>cd8tom;gfp*= 35 egg chambers, *tj>cd8tom;shgRNAi* = 37 egg chambers). **E** Quantification of coordination of nurse cell size changes during nurse cell dumping. The coe=icient of variance in nurse cell sizes within on egg chamber as a function of germline size. LOESS fitted with a 95% CI area. N egg chambers: (*tj>cd8tom;gfp*= 35 egg chambers, *tj>cd8tom;shgRNAi* = 37 egg chambers). **F** Quantification of percentage of stage 11 egg chambers with a blocked ring canal phenotype. N egg chambers: (*tj> cd8tom;gfp*= 13 egg chambers, *tj>cd8tom;shgRNAi* = 32 egg chambers). **G** Medial confocal section through egg chambers expressing *CD8::Tomato* under the control of TJ-Gal4 in follicle cells, stained for F-Actin and nuclear DNA (DAPI). Red arrow heads point to centripetal FCs that migrated between NCs and oocyte. White arrow heads point to actin cables that keep nuclei from blocking ring canals. Yellow arrowhead points to nurse cell nucleus pushed towards AFC-nurse cell interface. **H** Medial confocal section through egg chambers egg chambers expressing *CD8::Tomato* and *shg-RNAi* under the control of TJ-Gal4 in follicle cells, stained for F-Actin and and nuclear DNA (DAPI). Red arrows point to centripetal follicle cells that fail to migrate between nurse cells and oocyte. White arrows point nurse cell-oocyte interface lacking actin cables. Yellow arrows point to ring canals blocked by nurse cell nuclei.

### E-Cadherin spot junctions ensure nurse cell envelopment by AFCs

Simultaneously with nurse cell dumping, anterior follicle cells (AFCs) closely enwrap the nurse cells **(Fig. 4A**, **Fig. 5A, Fig. S4A)** and subsequently induce phagoptosis to remove nurse cell remnants (Timmons, Mondragon et al., 2017; Mondragon, Yalonetskaya et al., 2019; Serizier, Peterson et al., 2022). We sought to investigate whether the emerging E-Cadherin spot junctions play a role in the process of nurse cell envelopment by AFCs. To characterize this process, we analyzed the dynamics of envelopment during nurse cell dumping. Specifically, we quantified the proportion of the interface shared between nurse cells and AFCs as a function of nurse cell size **(Fig. 5B)**. Our analysis revealed that during the early stages of nurse cell size reduction, the proportion of the interface shared with AFCs remained stable. However, once nurse cells reached a critical size by half or more, the proportion of the surface covered by AFCs increased until the nurse cells were completely enveloped **(Fig. 5C, Fig. S4B)**. To investigate the role of spot junctions in this process, we combined data sets from egg chambers exhibiting the ring canal phenotype and those that did not, due to the strong penetrance of the phenotype **(Fig. 5D)**. In egg chambers expressing E-Cadherin RNAi in follicle cells, the increase in nurse cell envelopment by AFCs, when nurse cells shrank in size during dumping, was completely abolished **(Fig. 5E,F, Fig. S4B)**. Notably, in control nurse cells, the quantified interface proportion never dropped below 25%. In contrast, egg chambers lacking E-Cad spot junctions exhibited a significant population of nurse cells with interface proportions smaller than 25%. Strikingly, we also identified a substantial number of nurse cells that had completely detached from AFCs. **(Fig. 5D,F, Fig. S4B)**. These findings indicate that E-Cadherin junctions between AFCs and nurse cells are crucial for maintaining a stable contact surface during nurse cell dumping and enable the complete envelopment of nurse cells. Live imaging of egg chambers with labelled follicle cells during the dumping process further corroborated these results (**Fig. S4C,D**).

**Figure 5.**
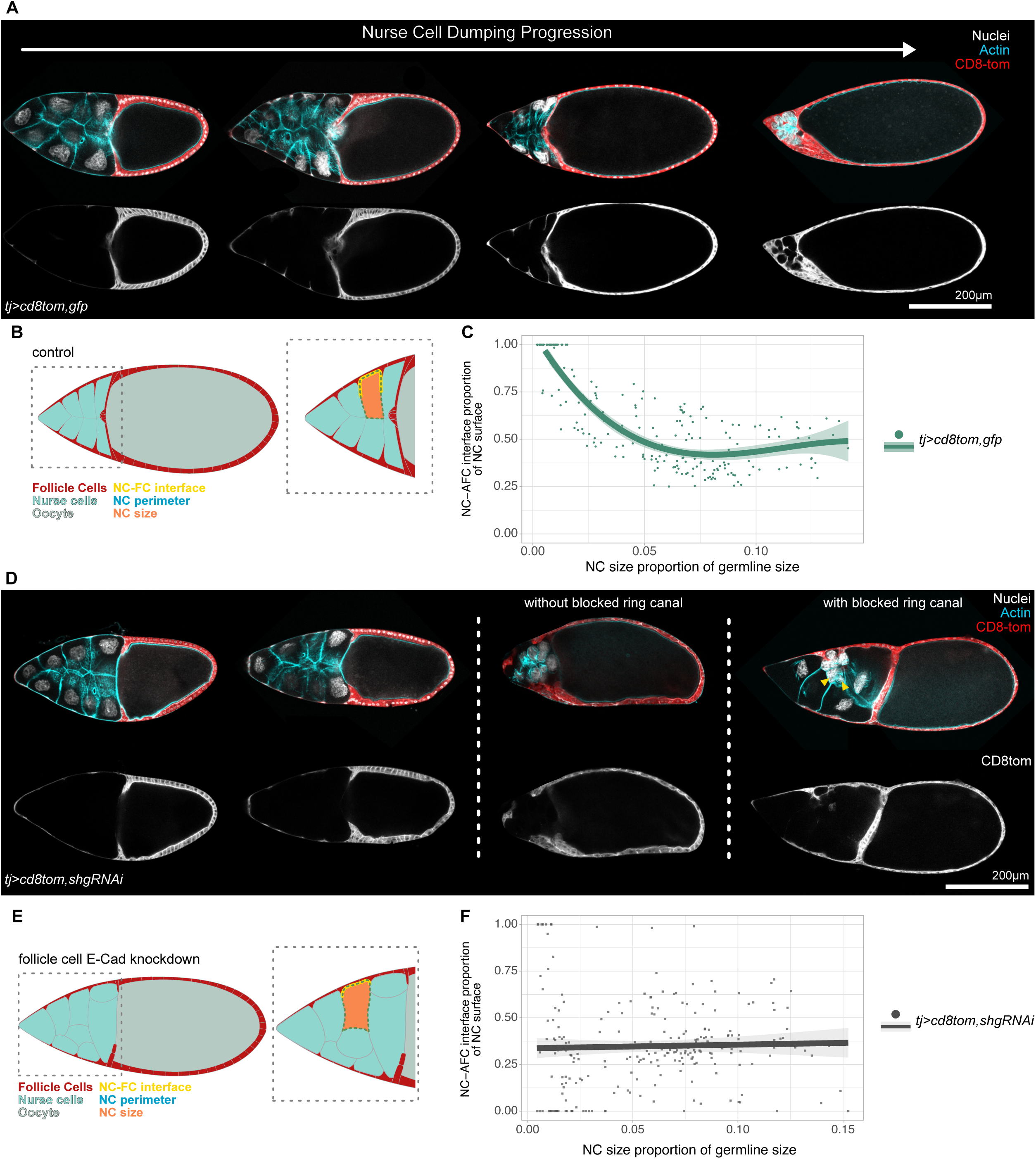
Spot junctions are required for AFC envelopment of nurse cells. **A** Confocal medial sections of egg chambers during nurse cell dumping, expressing *CD8::Tomato* and *GFP* under the control of TJ-Gal4 in follicle cells, and stained for nuclear DNA (DAPI) and F-Actin. **B** Illustration of control egg chambers during dumping. Quantification concept for nurse cell-AFC interface proportion is depicted. **C** Quantification of the nurse cell-AFC interface proportion of individual nurse cells as a function of nurse cell size (proportion of NC area of germline area). 3^rd^ order polynomial fit with 95%CI area. N egg chambers: (*tj>cd8tom,gfp* = 209 nurse cells from 30 egg chambers). **D** Confocal Medial sections of egg chambers during nurse cell dumping, expressing *CD8::Tomato* and *shg-RNAi (E-cadherin RNAi)* under the control of TJ-Gal4 in follicle cells, and stained for nuclear DNA (DAPI) and F-Actin. Two nurse cell dumping phenotypes are depicted. Yellow arrowheads point to nurse cells completely detached from follicle cells. **E** Illustration of egg chambers with follicle cell specific knockdown of E-cadherin during dumping. Quantification concept of nurse cells-AFC interface proportion is depicted. **F** Quantification of the nurse cells-AFC interface proportion of individual nurse cells as a function of nurse cell size (proportion of NC area of germline area). Linear fit with 95% CI area. N egg chambers: (*tj>cd8tom,shgRNAi* = 244 nurse cells from 37 egg chambers).

Taken together, these observations suggest that it is essential for AFCs to switch their E-Cadherin-based adhesion from intraepithelial contacts to the contact surface with the nurse cells to allow for the coordination between the two lineages in the intricate process of nurse cell dumping and enveloping.

## DISCUSSION

This study reveals a novel mechanism of cellular interaction during late *Drosophila* oogenesis by characterizing the formation and function of E-cadherin-based spot junctions at the interface between anterior follicle cells (AFCs) and nurse cells. These findings underscore the critical role of homophilic E-cadherin interactions in maintaining a stable soma-germline interface, which is essential for coordinating cellular behaviors and ensuring successful oocyte development.

Our results demonstrate that E-cadherin forms functional adhesive junctions at the apical surface of AFCs in contact with nurse cells, establishing a physical and regulatory interface between the two cell lineages. The redistribution of E-cadherin from intra-epithelial adherens junctions to apical spot junctions represents a striking example of epithelial plasticity. This process enables AFCs to adapt their adhesion profile to accommodate dynamic changes in their size and shape during oogenesis. Importantly, the dependency of spot junction formation on homophilic E-cadherin interactions highlights the specificity of this adhesive mechanism and its essential role in stabilizing soma-germline contacts. Our findings further suggest that the proportional increase of the apical surface area of AFCs acts as a spatial determinant for E-cadherin localization, where the increased probability of trans-bond engagement drives the stabilisation of E-cadherin at the apical surface. This model is supported by experiments showing ectopic formation of spot junctions in main body follicle cells (MBFCs) upon apical surface expansion, and is consistent with studies on E-cadherin stabilization in functional adhesion junction (Brasch, Harrison et al. 2012, Takeichi 2014). Our findings underscore the importance of cell geometry and contact area in guiding adhesion dynamics.

The stability and regulated organization of E-cadherin spot junctions during stages 10A to 11 is remarkable given the dramatic changes in size occurring in nurse cells during this period. Our quantitative analysis reveals that spot junction density, nearest-neighbour distribution, and organization remain constant despite significant increases in the interface area. This finding suggests a robust regulatory mechanism that ensures uniform adhesion strength and distribution, critical for maintaining tissue integrity and coordination during rapid morphological transitions. We furthermore observed that spot junctions localize preferentially to actin-free regions within the dense actin meshwork of nurse cells. Ultrastructural analysis revealed that this dense actin meshwork is made up by nurse cell microvilli extending into the intercellular space between nurse cells and AFCs. Furthermore, these microvilli are absent at regions of direct membrane contact, potentially corresponding to E-cadherin spot junctions. These observations highlight the interplay between adhesive and actin based structural features of the soma-germline interface, and future studies need to elucidate cross-talk during emergence of these distinct cellular structures. Moreover, we propose that spot junctions contribute significantly to the ability of AFCs to envelop the nurse cells at the final stages of dumping. As nurse cells rapidly shrink, the structural role of the junctions may be to stabilize and guide the dynamic interface between AFCs and the retreating nurse cell membranes. This stabilization could ensure the precise spatial positioning and morphological adaptation of AFCs as they close around the nurse cells. By facilitating this process, spot junctions may ensure efficient completion of nurse cell dumping while maintaining tissue integrity.

The apical localization of E-cadherin-based spot junctions in follicle cells represents an intriguing deviation from classical epithelial architecture, where apical surfaces typically face luminal spaces. From invertebrates to vertebrates this setup can be found during oogenesis, which suggests the need of a specialized adhesion mechanisms to accommodate the unique demands of oogenesis. Our findings add to the growing body of evidence that epithelial plasticity and apical-basal polarity diversification are central to the functional specialization of follicle cells in *Drosophila* and other species (Eckelbarger and Hodgson 2021, Spradling, Niu et al. 2022). These findings have broader implications for understanding adhesion-mediated communication and coordination in other developmental systems. The use of E-cadherin as a versatile adhesive molecule underscores its central role in diverse cellular contexts, from maintaining epithelial integrity to mediating inter-lineage interfaces in complex tissues. Future studies should aim to elucidate the signaling pathways that regulate E-cadherin redistribution and clustering. Additionally, exploring the mechanistic basis of cytoskeleton-adhesion integration at the soma-germline interface will provide further insights into how physical and biochemical cues are coordinated during tissue morphogenesis.

## MATERIALS AND METHODS

### *Drosophila* stocks and genetics

All experiments were performed on *Drosophila melanogaster*. Stocks (Supplementary Table 1) and experimental crosses (Supplementary Table 2) were maintained on standard fly food (10L water, 74.5g agar, 243g dry yeast, 580g corn meal, 552mL molasses, 20.7g Nipagin, 35mL propionic acid) at 18°C, 22°C and 25°C. Mosaic analysis was performed using the ‘flip-out’ and the mitotic FLP/FRT system (del Valle Rodriguez, Didiano et al. 2011). For follicle epithelium clones, *flp* expression was induced in young adult females using a heat shock varying from 4 to 20 min depending on the *hsflp* construct (*hsflp [122]* vs. *hsflp[1]*) for ‘flip-out’ experiments. Flies were fed yeast paste for 48 to 72 h prior to dissection.

### Immunohistochemistry and imaging

Ovaries were dissected and fixed in 4% paraformaldehyde/PBS for 15 min at 22°C. Washes were performed in PBS + 0.1% Triton X-100 (PBT). Samples were incubated with primary antibodies in PBT overnight at 4°C: mouse anti-β-catenin (1:100, DSHB, N27A1), rat anti-E-cadherin (1:50, DSHB, DCAD2), rabbit anti-GFP (1:200, Thermo Fisher, G10362), rat anti N-Cad (1:20, DSHB, DN-EXH8), mouse anti-PKC ζ (1:50, Santa Cruz, H-1,sc-17781). Ovaries were incubated with secondary antibodies for 2 h at 22 °C. DAPI (0.25 ng/μl, Sigma), Phalloidin (Alexa Fluor 488, Alexa Fluor 647 and Alexa Fluor 555, Molecular Probes, or Phalloidin-TRITC, Sigma) was used to visualize DNA and filamentous Actin. Following secondary antibodies were used: goat anti-mouse Alexa Fluor488 (Abcam, AB150117, 1:500), goat anti-rat Alexa Fluor488 (Abcam, AB150153, 1:500), goat anti-rabbit Alexa Fluor488 (Invitrogen, A11008, 1:500), donkey anti-mouse Alexa Fluor555 (Abcam, AB150110, 1:500), donkey anti-rat Alexa Fluor555 (Abcam, AB150154, 1:500), donkey anti-mouse Alexa Fluor647 (Abcam, AB150111, 1:500), donkey anti-rat Alexa Fluor647 (Abcam, AB150155, 1:500), goat anti-rabbit Alexa Fluor647 (Invitrogen, A21244, 1:500). Samples were mounted using Molecular Probes Antifade Reagents and imaged using Leica TCS SP8 confocal microscopes. Control and experimental samples were processed in parallel, and images were acquired using the same confocal settings.

Nurse cell dumping was imaged in the following medium: 7.6ml Schneiders Medium, 2ml FBS, 0.4mg/ml Insulin, pH=6.9-7.

### Image Acquisition, Analysis and Quantification

Images were obtained with a LEICA TCS SP8 using the software LAS X for confocal imaging, ZEISS LSM 880 Examiner for Airyscan imaging, and Axio Oberserver 7 for Live Imaging of nurse cell dumping. Images were processed in FIJI (Schindelin, Arganda-Carreras et al. 2012). Statistical analysis and generations of graphs were performed in R (R version 4.0.5), GraphPad (GraphPad Prism 9).

### Spot Junction Distribution Quantification

Select right plane of Airyscan Images of Spot Junctions, up to 3 planes as Max. Proj.. E-cadherin and b-cat channels were merged to create a spot junction channel. Measurement area was generated manually to exclude areas out of focus and sSpot junction channel was thresholded with Triangle or Moments method. The despeckle function was applied. Analyze particles function with 0.05µm^2^ – Infinity was applied. Measurements included centroids and area. ROIs were added to ROI manager. NND were measured. All ROIs were combined into one ROI with the combine function and then saved as spot junctions ROI. Mask was generated from Spot Junctions ROI and saved as spot junctions mask. 3D Object Counter without numbering was applied. Output was directly used for 3D shuffle. Original spot junction mask was shuffled 5 times with applied measurement area and saved each time (Fig. S2). Shuffled images were binarized. Then same procedure as for the original image was performed to extract coordinates, NNDs and sizes.

### Germline size, nurse cell size and nurse cell-follicle cell interface area quantification

Egg chamber morphology measurements were performed in FIJI, using the polygon, line tool. Germline area was determined in 2D medial cross sections (through the anterior and posterior pole (Ref Weichselberger 2022). NC-FC interface was calculated by approximating the nurse cell compartment to a cone and calculating the lateral surface area of a cone with the measured radius and slant height from a 2D medial section of the egg chamber. Individual nurse cell sizes were quantified as areas in confocal sections through the center of the nurse cell. Wild type and control egg chambers were staged by previously described criteria (Koch 1963, Horne-Badovinac and Bilder 2005).

### Electron microscopy

Isolated ovarioles were incubated in Schneider’s medium containing 2.5% glutaraldehyde and 2% paraformaldehyde for 30 min, followed by incubation in PHEM buffer with fixatives for additional 30 min. Samples were flat embedded in 2% low melting agarose and placed at 4 C° for 10 minutes until polymerization. Blocks of agarose were transferred to PHEM buffer with fixatives at 4°C overnight. Post-fixation, embedding, and imaging were performed as described in (Schuessele, Hoernstein et al. 2016).

## ACKNOWLEDGEMENTS

We thank the sta_ of the Life Imaging Center (LIC) in the Hilde Mangold House (HMH) of the University of Freiburg for help with their confocal microscopy resources, and the excellent support in image recording. We specifically thank the DFG for supporting our imaging work through project number 414136422. We thank the Bloomington *Drosophila* Stock Center (BDSC), the Vienna *Drosophila* Stock Collection (VDRC) and the Developmental Studies Hybridoma Bank (DSHB) for providing fly stocks and antibodies. We thank David Bilder, Katja Röper, Sally Horne-Badovinac, Kim McCall, Thomas Lecuit, Ulrich Tepass for sharing reagents. We also thank the SGBM and IMPRS graduate schools for supporting our doctoral researchers.

## AUTHOR CONTRIBUTIONS

Conceptualization VW, AKC

Experimental Validation VW, RB

Experimental Investigation VW, MRF

Writing VW, AKC

Visualization VW, AKC

Supervision AKC

Funding Acquisition AKC

---

**Supplementary Table 1.**
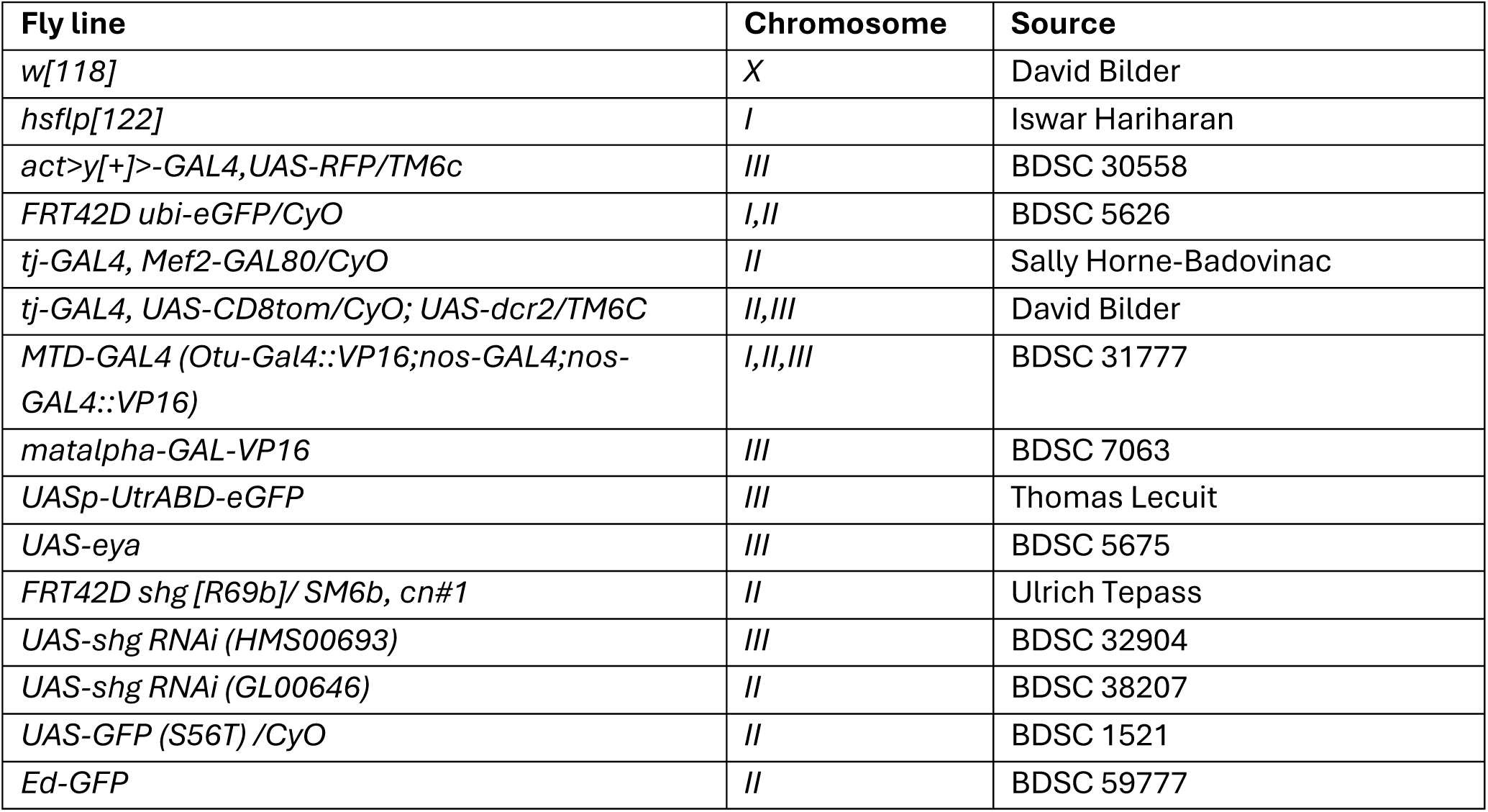
– Fly strains.

**Supplementary Table 2.**
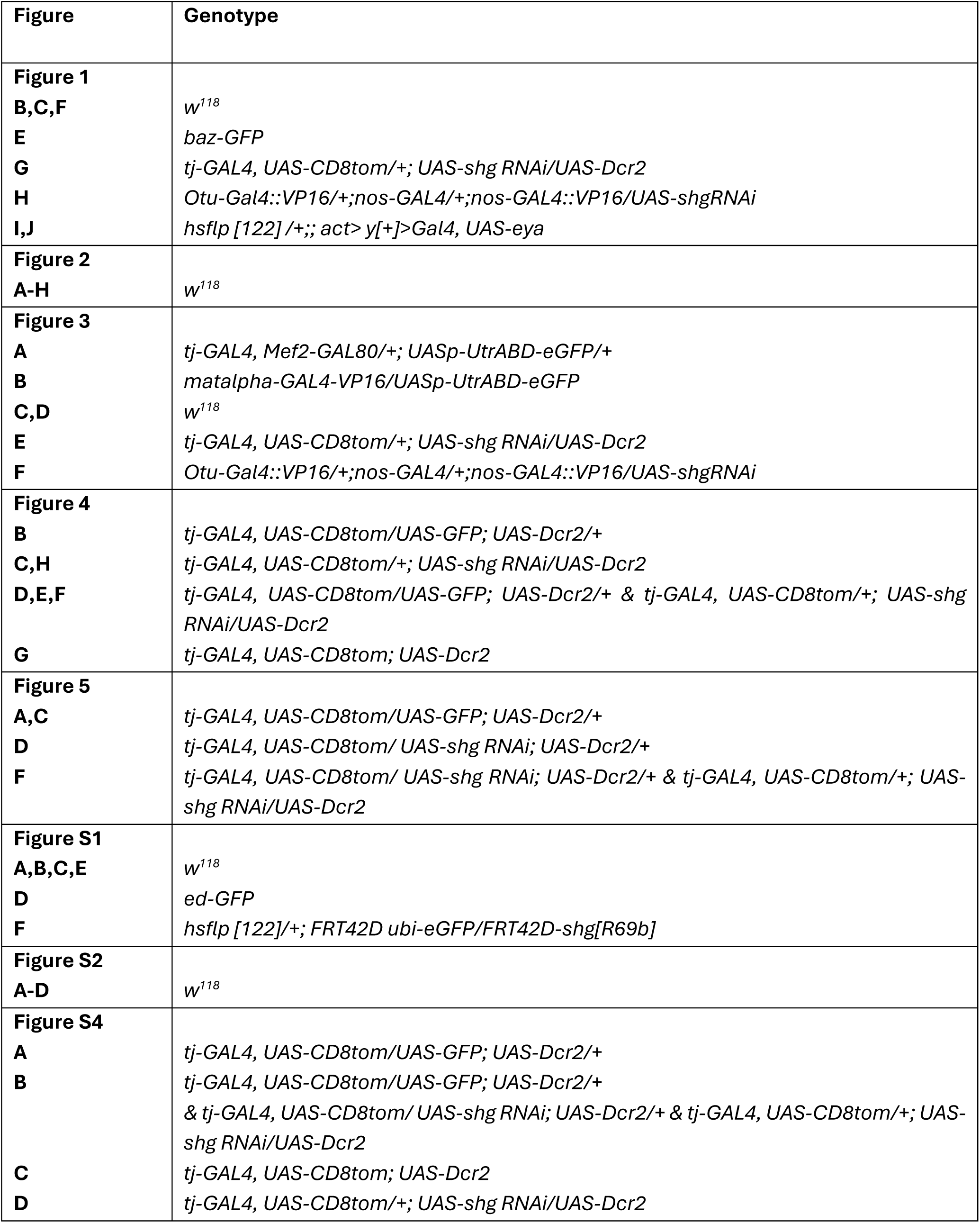
– Experimental genotypes.

**Figure S1.**
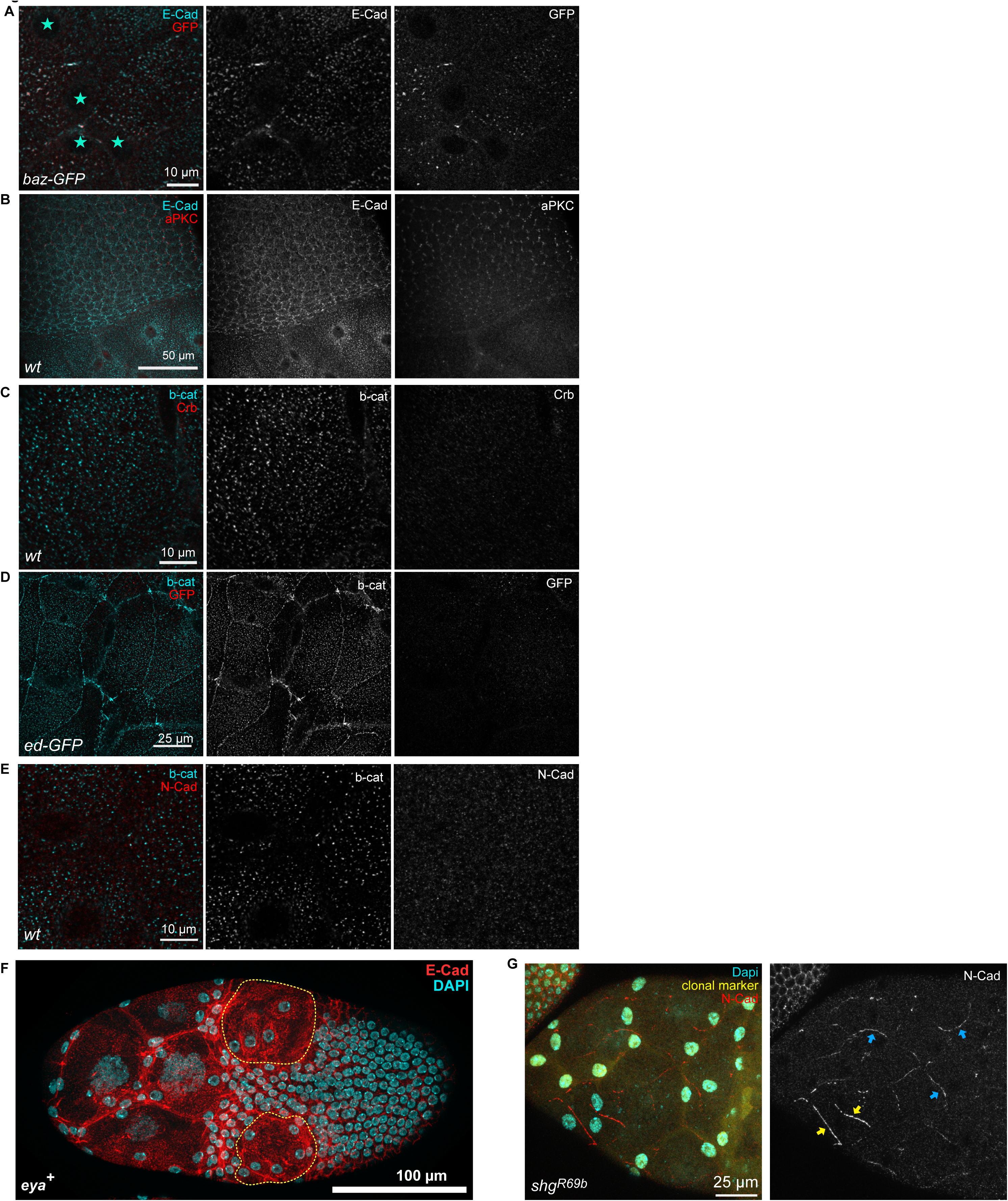
Spot junction emergence. **A** Max. projection of confocal images depicting AFCs in stage 10A egg chamber, expressing *bazooka(baz)-GFP* and stained for E-cadherin and GFP. **B** Max. projection of confocal images of a stage 10A wt egg chamber, depicting AFCs and follicle cells in contact with the oocyte, stained for E-cadherin and aPKC. **C** Max. projection of confocal images of a stage 10A wt egg chamber depicting AFCs stained for β-catenin and Crumbs (Crb). **D** Max. projection of confocal images depicting AFCs of a stage 10A egg chamber expressing *echinoid*(*ed)-GFP*, stained for β-catenin and GFP. **E** Max. projection of confocal images of a stage 10A wt egg chamber depicting AFCs stained for E-cadherin and N-Cad. **F** Max. projection of confocal images depicting egg chamber with clones ectopically expressing eya, stained for E-cad and Nuclei (DAPI). Yellow dotted line encircles cells with ectopic expression of eya. Note lack of E-cadherin at follicle cell junctions and presence of E-cadherin spots on their apical surface. **G** Max. projection of a stage 10A egg chamber with clones homozygous mutant for the *E-cadherin shg^R69b^* allele (clonal marker negative), stained for nuclear DNA (DAPI) and N-Cadherin. Blue arrows mark intra-epithelial junctions between control cells, yellow arrows point to intra-epithelial junctions lacking E-cadherin.

**Figure S2.**
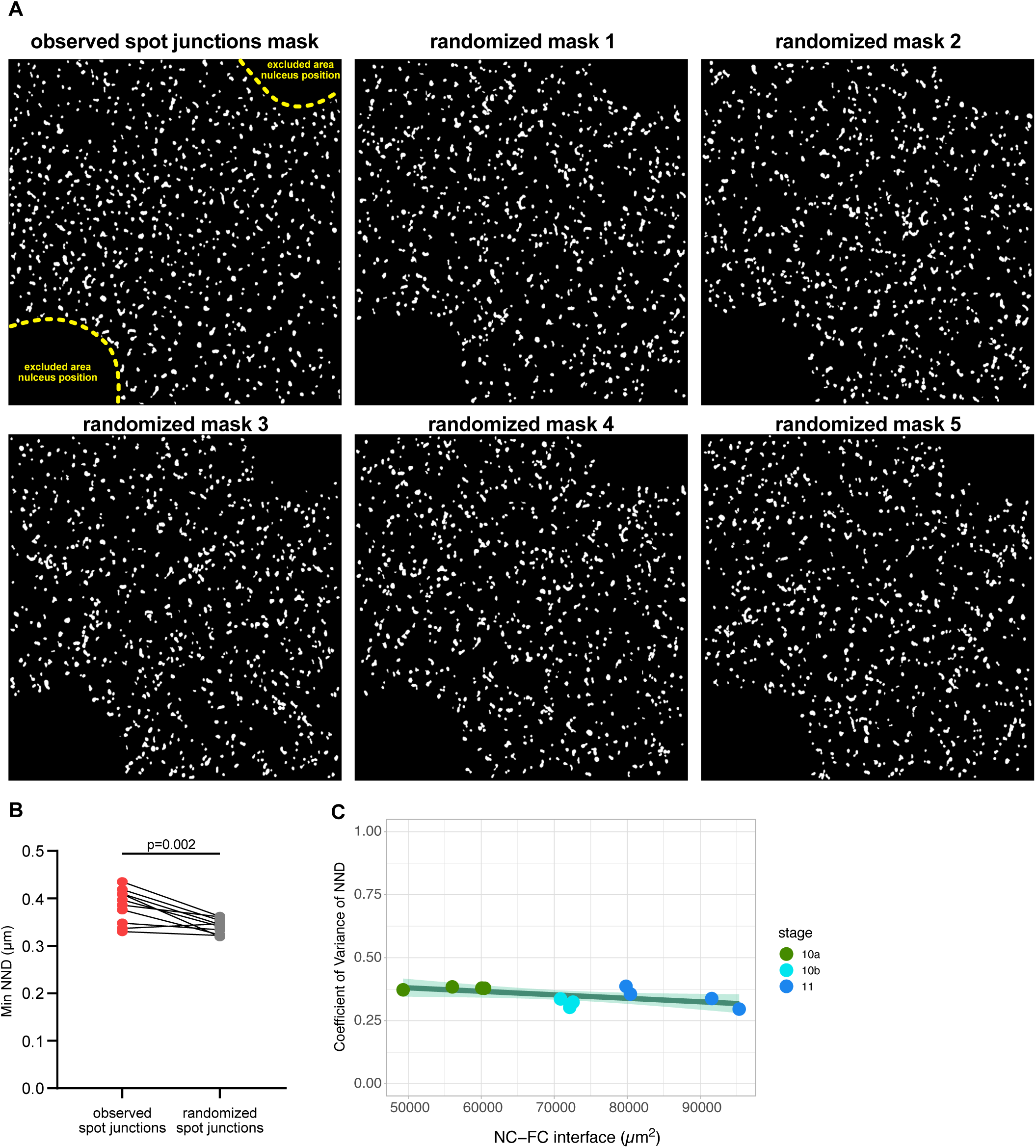
Spot junction distribution. **A** Example of a segmented masks for spot junctions, generated from Airyscan confocal images, and the respective 5 randomized distributions used for analysis (see Materials&Methods). **B** Two-tailed paired t-test of the minimal Nearest Neighbor Distance (Min NND) between E-cadherin spot junctions. Observed spot junction organization vs randomized organization (see Fig. S2A & Methods). n=11 egg chambers. **C** Coe=icient of Variance (CV) of Nearest Neighbor Distances (NNDs) as a function of nurse cell-to-follicle cell interface size of wt egg chambers. Linear fit with a 95% CI area. n=11 egg chambers.

**Figure S3.**
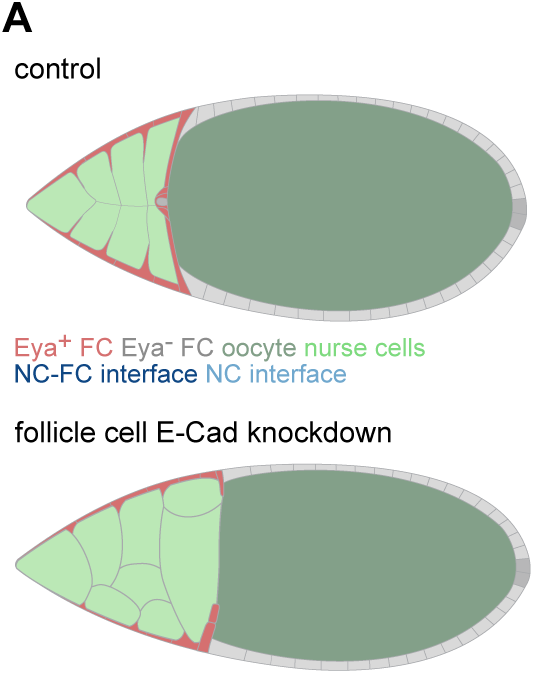
Spot junction function during nurse cell dumping. **A** Illustration of a stage 11 wt egg chamber, and a stage 11 egg chamber expressing *shg-RNAi (E-cadherin RNAi)* to knockdown E-Cadherin in follicle cells.

**Figure S4.**
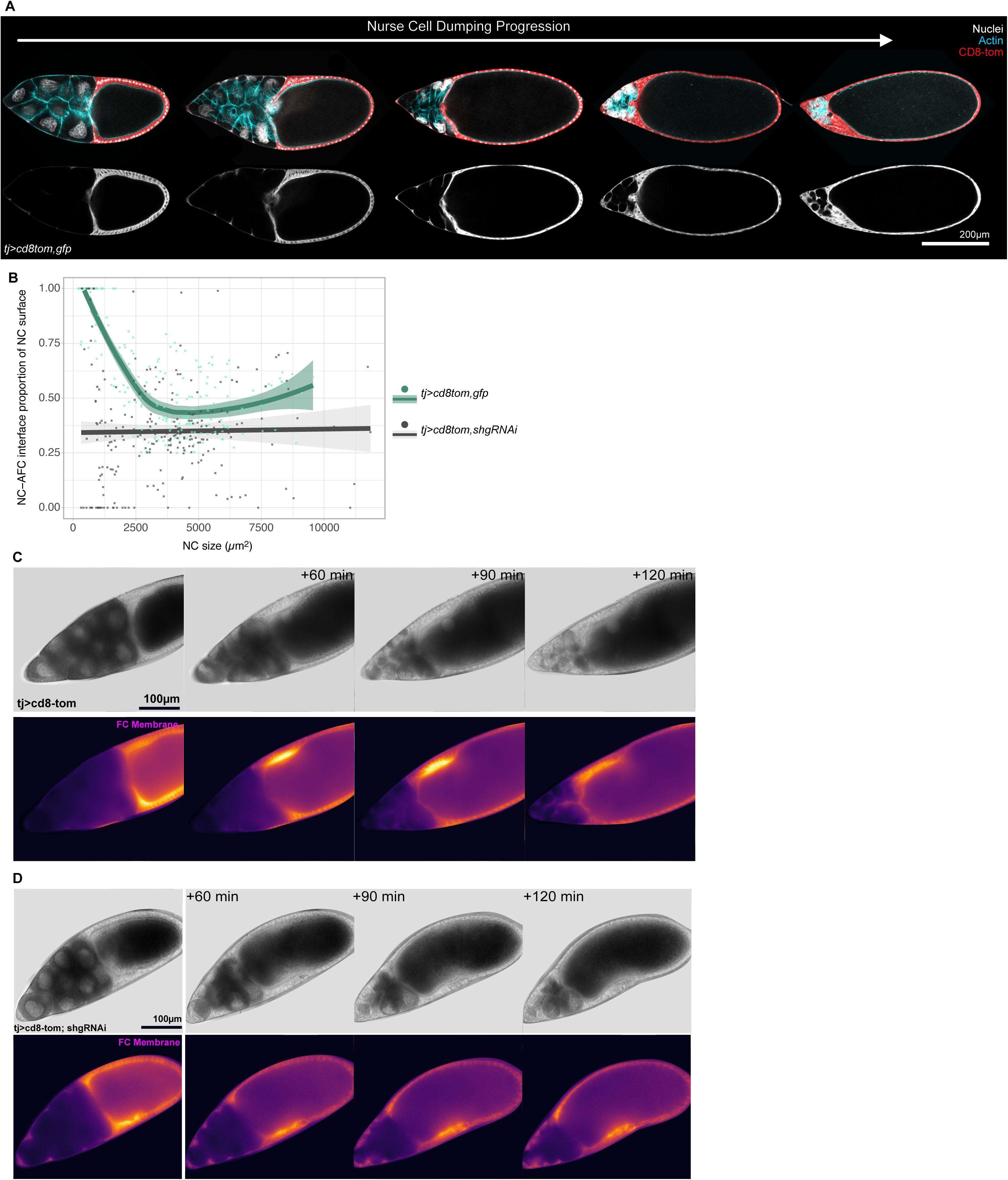
Spot junctions are essential for nurse cell envelopment during dumping. **A** Confocal medial sections of egg chambers during nurse cell dumping, expressing *CD8::Tomato* and *GFP* under the control of TJ-Gal4 in follicle cells, and stained for nuclear DNA (DAPI) and F-Actin. **B** Quantification of the nurse cell-AFC interface proportion of individual nurse cells as a function of nurse cell size. *tj>cd8tom,gfp* LOESS fitted with mean and 95%CI area. (*tj>cd8tom,gfp* = 209 nurse cells from 30 egg chambers). *tj>cd8tom,shgRNAi* linear fit with 95% CI area. (*tj>cd8tom,shgRNAi* = 244 nurse cells from 37 egg chambers). **C** Images of a live movie of an egg chamber expressing *CD8::Tomato* and *GFP* under the control of TJ-Gal4 in follicle cells during nurse cell dumping. Brightfield and *CD8::Tomato* signals are shown. Note how follicle cells (*CD8-Tom*) envelop nurse cells. **D** Images of a live movie of an egg chamber expressing *CD8::Tomato* and *shg-RNAi (E-cadherin RNAi)* under the control of TJ-Gal4 in follicle cells during nurse cell dumping. Brightfield and *CD8::Tomato* signals are shown. Note how follicle cells (CD8-tom) fail to envelop nurse cells.

**Movie S1 Nurse cell dumping in wild type egg chambers**

Live imaging of a wild type chamber expressing *CD8::Tomato* and *GFP* under the control of TJ-Gal4 in follicle cells during nurse cell dumping. Brightfield (left) and *CD8::Tomato* (right) signals are shown. Note how follicle cells (*CD8-Tom*) envelop nurse cells.

**Movie S2 Nurse cell dumping in egg chambers expressing E-cadherin RNAi**

Live imaging of a wild type chamber expressing *CD8::Tomato* and *shg-RNAi (E-cadherin RNAi)* under the control of TJ-Gal4 in follicle cells during nurse cell dumping. Brightfield (left) and *CD8::Tomato* (right) signals are shown. Note how follicle cells (CD8-tom) fail to envelop nurse cells.

